# Separating Measured Genetic and Environmental Effects: Evidence Linking Parental Genotype and Adopted Child Outcomes

**DOI:** 10.1101/698464

**Authors:** Benjamin W. Domingue, Jason Fletcher

**Affiliations:** Graduate school of Education, Stanford University; La Follette School of Public Affairs, Department of Sociology, and Center for Demography of Health and Aging, University of Wisconsin-Madison

**Keywords:** Indirect genetic effects, Polygenic Score, Educational Attainment, Wisconsin Longitudinal Study, Adoption Study

## Abstract

There has been widespread adoption of genome wide summary scores (polygenic scores) as tools for studying the importance of genetics and associated lifecourse mechanisms across a range of demographic and socioeconomic outcomes. However, an often unacknowledged issue with these studies is that parental genetics impact both child environments and child genetics, leaving the effects of polygenic scores difficult to interpret. This paper uses multi-generational data containing polygenic scores for parents (n=7,193) and educational outcomes for adopted (n=855) and biological (n=20,939) children, many raised in the same families, which allows us to separate the influence of parental polygenic scores on children outcomes between environmental (adopted children) and environmental and genetic (biological children) effects. Our results complement recent work on “genetic nurture” by showing associations of parental polygenic scores with adopted children’s schooling, providing additional evidence that polygenic scores combine genetic and environmental influences and that research designs are needed to separate these estimated impacts.

## 1. Introduction

This paper focuses on the issue of separating genetic and environmental influences in important adult socioeconomic outcomes. The rapid advancement in the collection, measurement, and understanding of genetic data has led to new genetic discoveries linking specific genetic variants with health outcomes as well as schooling outcomes (1–3). While, individually, these variants do not have large effects (4), the development of genome wide summary scores (polygenic scores, PGS) offers the possibility of a genetically-based analytic tool that is predictive (5) of important adult outcomes, including health (6) and disease outcomes (7) but also behavioral outcomes such as schooling (8).

Many studies have begun to examine the characteristics of a PGS constructed to predict years of schooling (i.e., educational attainment). Studies have examined associations between the educational attainment PGS and outcomes of interest at different stages of the lifecourse including childhood and adolescence (9–11), middle age (9,12,13), and later-life (14,15). Other studies have examined the impact of the educational attainment PGS on economic attainments such as earnings and wealth (16–18). Still others have used this PGS to examine gene-environment interactions (19–22). While these studies have begun to offer novel insights into human behavior, they also raise questions about how to interpret associations between this polygenic score and the various outcomes in question.

The issue of how to interpret polygenic score associations with phenotypes has been repeatedly raised by scholars, especially when it comes to educational attainment (23). PGS are created by summing up results of GWAS effect estimates across the genome; however, the acknowledgement that GWAS, as a research design, is not adequate to separate causal and non-causal genetic variants when estimating effects leads to the implication that PGS measures capture both causal and non-causal genetic effects on the phenotype.

As a first exploration of this issue, researchers have examined the associations between PGS and phenotype with a sibling design—leveraging differences in PGS between biological siblings as a means to attempt to separate causal and non-causal components of the PGS. Specifically, analysis of siblings is powerful as it minimizes the possibility for confounding due to factors that do not vary within-family. Early evidence (19,23,24) suggested that the PGS for educational attainment is indeed capturing a crucial signal of individual difference in the sense that the sibling with the higher PGS tended to also have more schooling. However, more recent work based on siblings suggests that associations between PGS and educational attainment are substantially smaller when drawn from family-based samples as compared to association estimates drawn from unrelated individuals (25). This suggest the possibility of bias—in particular, upwards bias in PGS estimates—when non-family samples are analyzed (26).

Relatedly, newer work has begun to produce evidence of “social genetic effects”, where the genetics of a focal individual’s social connections could play a role in the focal individual’s outcomes (27–29). Social genetic effects may, in many cases, be phenotypic effects in action— they may be mediated by observable phenotypes of an individual’s social peers—but, to the extent that they exist, they may lead to conceptual and methodological difficulties by adding a new layer of confounding to the PGS. The effects of polygenic scores would then need to be interpreted accordingly (e.g., they capture processes that are not within-person) and even sibling difference models may suffer from stable unit treatment value assumption (SUTVA) violations that can produce results that are biased in either direction (30).

A particularly important potential source of confounding with respect to the educational attainment polygenic score is the observation that variation in the parental genotype associated with educational attainment may also be associated with differences in the environments that parents create for their children. Were this hypothesis true, parental genetics would have implications for both child environments and child genetics, offering an additional reason the estimated effects of polygenic scores would be difficult to interpret as purely within-person genetic effects. Indeed, emerging work is beginning to examine the impacts of parental genotype (either net of child genotype or focusing on the non-transmitted portion) on children’s outcomes, termed “genetic nurture” (31–35). Note that while our focus on educational attainment is motivated primarily by its centrality to many social science questions, the findings of Kong et al. (31) suggest that educational attainment may be a phenotype that is especially impacted by genetic nurture.

Existing estimates of genetic nurture may understate the total contribution of family background to child’s phenotype. Kong et al. (31) partitions the parental genetic effects on offspring phenotype into transmitted and non-transmitted alleles. A PGS created from the non-transmitted alleles are the “indirect genetic effects” (or genetic nurture) and a PGS created from the transmitted alleles are the “direct genetic effects” in this framework. However, this partitioning exercise does not account for the fact that the transmitted alleles can affect children both directly and indirectly through parentally provided environments. In this paper, we introduce an alternative assessment that does not partition between transmitted and non-transmitted allele and instead estimates the “genetic nurture” effect via a different design.

This paper offers novel evidence on the magnitude of genetic nurture effects by using multi-generational data that collected polygenic scores from 7,200 parents and outcomes for 22,000 of their adopted and biological children, which allows us to separate the influence of parental polygenic scores on children outcomes between environmental (i.e. genetic nurture) (adopted children) and environmental and genetic (biological children) effects. We do this by focusing on a test of whether the polygenic score of a parent predicts the educational attainment of adopted offspring (to whom they have not transmitted their genes).

## 2. Methods

### 2A. Data

The Wisconsin Longitudinal Study (WLS) is a long-term study of over 10,000 men and women based on a random 1/3 sample of the graduating high school class of 1957 in Wisconsin (36). These individuals continue to be followed through the present. Alongside the respondent who graduated in 1957, siblings were also impaneled (we refer to “graduate” and “sibling” respondents throughout to distinguish between the two types). The WLS provides an opportunity to study the life course, relationships, family functioning, physical and mental health and well-being, and morbidity and mortality from late adolescence through 2008. WLS data also cover social background, youthful aspirations, schooling, military service, labor market experiences, family characteristics and events, social participation, psychological characteristics, and retirement. Further, in the latest complete round of survey collection, the respondents have provided biological specimens for DNA analysis. The WLS now contains genome-wide DNA information for many of their respondents.

The WLS also asks respondents about their offspring. In addition to the graduate and sibling respondents reporting information on their own lives, they also report information on all their surviving children, including both the relationships with each child and information about that child’s education. For each child, the respondent was asked their relationship with that child (possible choices: biological, adopted, step, foster, legal ward, niece/nephew, other non-relative, child of lover/partner). Given complexities associated with these different relationships (e.g., assortative mating may induce correlations between polygenic scores of parents and children of their partners, environments may vary across these different familial relationships), we focus on adopted (n=855) and biological children (n=20,939) in remaining analyses.

In total, we observe 21,794 parent/child pairs for 7,193 WLS parents who have been genotyped. We note two points about this sample. First, the WLS parents are clustered into 5,695 families (1493 families have two WLS parents and two families contain three and four WLS parents respectively). This clustering has implications for statistical inference as parents within the same family are not independent (e.g., they grew up in a similar environment); we cluster standard errors accordingly as discussed below. Second, our analytic sample (N=7,193 WLS parents) shows small differences on key demographics from the full WLS sample (N=19,046). Focusing on the graduate respondents in each sample, graduates in the analytic sample were slightly younger (they were born about a week later; the difference was significant, p=0.009) and slightly less likely to be male (47% of our graduates were male as compared to 48% in the full sample; the difference was not significant). However, our analytic sample had a higher concentration of graduates; in the full WLS sample, 54% of the respondents are graduates while in our analytic sample 65% of the respondents are graduates.

### 2B. Measures

#### Parent Polygenic Score

We use the educational attainment polygenic score constructed by WLS and the SSGAC following the completion of the third generation educational attainment GWAS (8). Complete details regarding score construction are available elsewhere (37); we offer a summary of key points here. As WLS respondents were used in the main educational attainment GWAS, GWAS estimates were re-computed without the WLS respondents. These estimates were then used in combination with LDpred (38) to create a polygenic score. Given issues associated with polygenic prediction in diverse samples (39,40), scores were constructed only for respondents of European ancestry. We residualize the score based on the first ten principal components and then standardized to have mean=0 and SD=1 in this ancestrally homogeneous subsample (N=8,509).

#### Parent Education

Parental education is compiled via the collection of responses given by WLS respondents to education-related questions in the various waves of WLS. In particular, as in both the original GWAS (8) and in other work with this data (18,20), we use the maximum number of years of education reported by a WLS respondent over the different waves of data collection.

#### Child education

Offspring educational attainment was collected in 2004 when WLS respondents were asked versions of the question “What is the highest grade of regular school your child ever attended?” for all of their children. We used the coding of the WLS variable (e.g., https://www.ssc.wisc.edu/wlsresearch/documentation/waves/?wave=grad2k&module=gkid) which was meant to capture the years of schooling (e.g., a high school graduate is coded as having 12 years of education; note that we coded those with GEDs as having 11 years of education).

Table 1 presents descriptive statistics for the analytic sample and stratifies these descriptive statistics by child status (biological or adopted). Nearly two thirds of the children have parents who are graduate respondents (as opposed to siblings of WLS graduates). The graduate respondents were born in a narrow window around 1939 (recall that they are the graduating class of 1957); there is substantially more variation in the birthyears of the sibling respondents (SD for graduates=0.52, SD for siblings=6.8). By construction, the education polygenic score of parents is centered at 0; however, note that the polygenic scores of parents who have adopted children are higher than those who have biological children. Parents of adopted children also have higher educational attainments. The birth years of the children are centered in the mid-1960s although, as expected, the birth years of adopted children are later. Children’s educational attainment is over 14 years on average, although this figure is lower for adopted children.

**Table 1.** Summary Statistics: Wisconsin Longitudinal Study; Comparing Biological and Adopted Children

A. Child-Level Descriptives (Means and Standard Deviations, SD)
|  | All | Biological | Adopted |
| --- | --- | --- | --- |
| % Graduates | 0.66 | 0.66 | 0.67 |
| SD | 0.47 | 0.47 | 0.47 |
| Parent Birthyear | 1939.03 | 1939.01 | 1939.55 |
| SD | 3.94 | 3.95 | 3.59 |
| Parent PGS | -0.01 | -0.01 | 0.13 |
| SD | 0.99 | 0.99 | 0.95 |
| Parent Edu | 13.61 | 13.57 | 14.39 |
| SD | 2.31 | 2.29 | 2.58 |
| Child Birthyear | 66.86 | 66.71 | 70.62 |
| SD | 6.46 | 6.40 | 6.87 |
| Child Edu | 14.40 | 14.43 | 13.74 |
| SD | 2.42 | 2.41 | 2.65 |
| N | 21794 | 20939 | 855 |

B. Parent-level Descriptives (Means and Standard Deviations, SD)
|  | All | No Adopted | Adopted |
| --- | --- | --- | --- |
| % Graduates | 0.65 | 0.65 | 0.67 |
| SD | 0.48 | 0.48 | 0.47 |
| Parent Birthyear | 1939.41 | 1939.4 | 1939.6 |
| SD | 4.11 | 4.14 | 3.71 |
| Parent PGS | 0 | -0.01 | 0.11 |
| SD | 1 | 1 | 0.96 |
| Parent Education | 13.75 | 13.7 | 14.35 |
| SD | 2.36 | 2.33 | 2.59 |
| N | 7193 | 6648 | 545 |

### 2C. Analysis

So as to evaluate differential prediction of the PGS between biological and adopted children, we focus on linear models that contain interactions between the parental PGS and the child status (biological or adopted). For child j of parent i, we focus on linear models of the form

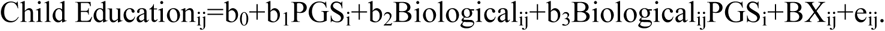

Our focal coefficient is b_1_ which captures the association of parent PGS with adopted offspring educational attainment. Our key hypothesis is that b_1_>0, indicating that the parental PGS predicts the educational attainment of their adopted children. Coefficients b_2_ and b_3_ describe differences between biological and adopted children. The b_2_ coefficient describes differences in schooling between biological and adopted children that may occur if there are systematic differences in environments associated with adoption status (e.g., parents construct different rearing environments for biological versus adopted children, the challenges faced by adoptees predating placement for adoption may have effects on their later attainments, etc.). Given the fact that biological children tend to have more years of education than adopted children, we anticipate b_2_>0. The b_3_ coefficient, which we expect to be positive given that it will also capture part of the effect of the genetics of biological children directly on their phenotype, will inform us about differences in the association between parental PGS and offspring educational attainment for biological versus adopted siblings. X is a matrix of controls including child sex and age (birth year); previous work (41) has shown the need for interactions in such models and we include those terms as needed. Standard errors are clustered—in particular, we use sandwich techniques (42) to obtain cluster-robust standard errors (43)—based on the family structuring of WLS participants (graduates plus siblings).

## 3. Results

Amongst biological children, the parental PGS is a robust predictor of child attainment. A one SD increase in the parental PGS predicts nearly an extra half year of schooling for a biological child (Table 2; M3). This is consistent with a large amount of previous work using a similar polygenic score (18,20,28). We next considered a test of the basic question: does the parental PGS predict educational attainment for non-biological offspring in the form of adopted children? In a baseline analysis, we considered a model (M0) containing main effects and interactions between parent PGS and child status. While the parental PGS is more predictive of the educational attainment of biological offspring, it also predicts variation in the variation of educational attainment in adopted children. A one standard deviation increase in PGS is associated with roughly an extra quarter year of schooling in adopted children (b=0.253, p=0.025). Results are summarized in Figure 1; the fact that the parent PGS is more predictive of educational attainments for biological children (i.e., b_3_>0) but that it also predicts variation in outcomes for adopted children (i.e., b_1_>0) is readily observed. Results were similar when we added controls for birthyear. We obtained a slightly attenuated result when we consider a model saturated with main effects and interactions of child gender (M2). We also obtained comparable results when we focus on models stratified by child status (M3 & M4).

**Table 2.** Associations between child educational attainment and parental polygenic score (PGS) for educational attainment.

|  | M0 | M1 | M2 | M3-bio<br>only | M4-adopted<br>only |
| --- | --- | --- | --- | --- | --- |
| Intercept | 13.708 | 14.097 | 14.147 | 14.407 | 14.140 |
| SE | 0.105 | 0.110 | 0.154 | 0.032 | 0.154 |
| p-value | <1E-5 | <1E-5 | <1E-5 | <1E-5 | <1E-5 |
| PGS | 0.253 | 0.263 | 0.227 | 0.479 | 0.273 |
| SE | 0.113 | 0.106 | 0.109 | 0.022 | 0.108 |
| p-value | 2.53E-02 | 1.36E-02 | 3.73E-02 | <1E-5 | 1.16E-02 |
| Bio Child | 0.728 | 0.341 | 0.258 |  |  |
| SE | 0.107 | 0.112 | 0.157 |  |  |
| p-value | <1E-5 | 2.37E-03 | 9.94E-02 |  |  |
| PGS:Bio Child | 0.227 | 0.216 | 0.224 |  |  |
| SE | 0.115 | 0.109 | 0.109 |  |  |
| p-value | 4.72E-02 | 4.73E-02 | 4.00E-02 |  |  |
| Child birthyear |  | -0.098 | -0.098 | 0.006 | -0.097 |
| SE |  | 0.023 | 0.023 | 0.005 | 0.023 |
| p-value |  | 1.99E-05 | 2.64E-05 | 2.23E-01 | 3.35E-05 |
| Bio Child:Child birthyear |  | 0.105 | 0.104 |  |  |
| SE |  | 0.023 | 0.024 |  |  |
| p-value |  | <1E-5 | 1.23E-05 |  |  |
| PGS:Child birthyear |  | 0.003 | 0.002 |  |  |
| SE |  | 0.004 | 0.004 |  |  |
| p-value |  | 4.90E-01 | 6.18E-01 |  |  |
| Male Child |  |  | -0.086 | 0.070 | -0.075 |
| SE |  |  | 0.207 | 0.047 | 0.207 |
| p-value |  |  | 6.76E-01 | 1.32E-01 | 7.15E-01 |
| Bio Child:Male Child |  |  | 0.157 |  |  |
| SE |  |  | 0.210 |  |  |
| p-value |  |  | 4.55E-01 |  |  |
| PGS:Male Child |  |  | 0.064 |  |  |
| SE |  |  | 0.044 |  |  |
| p-value |  |  | 1.45E-01 |  |  |
| N | 21021 | 21010 | 21010 | 20189 | 821 |

**Figure 1.**
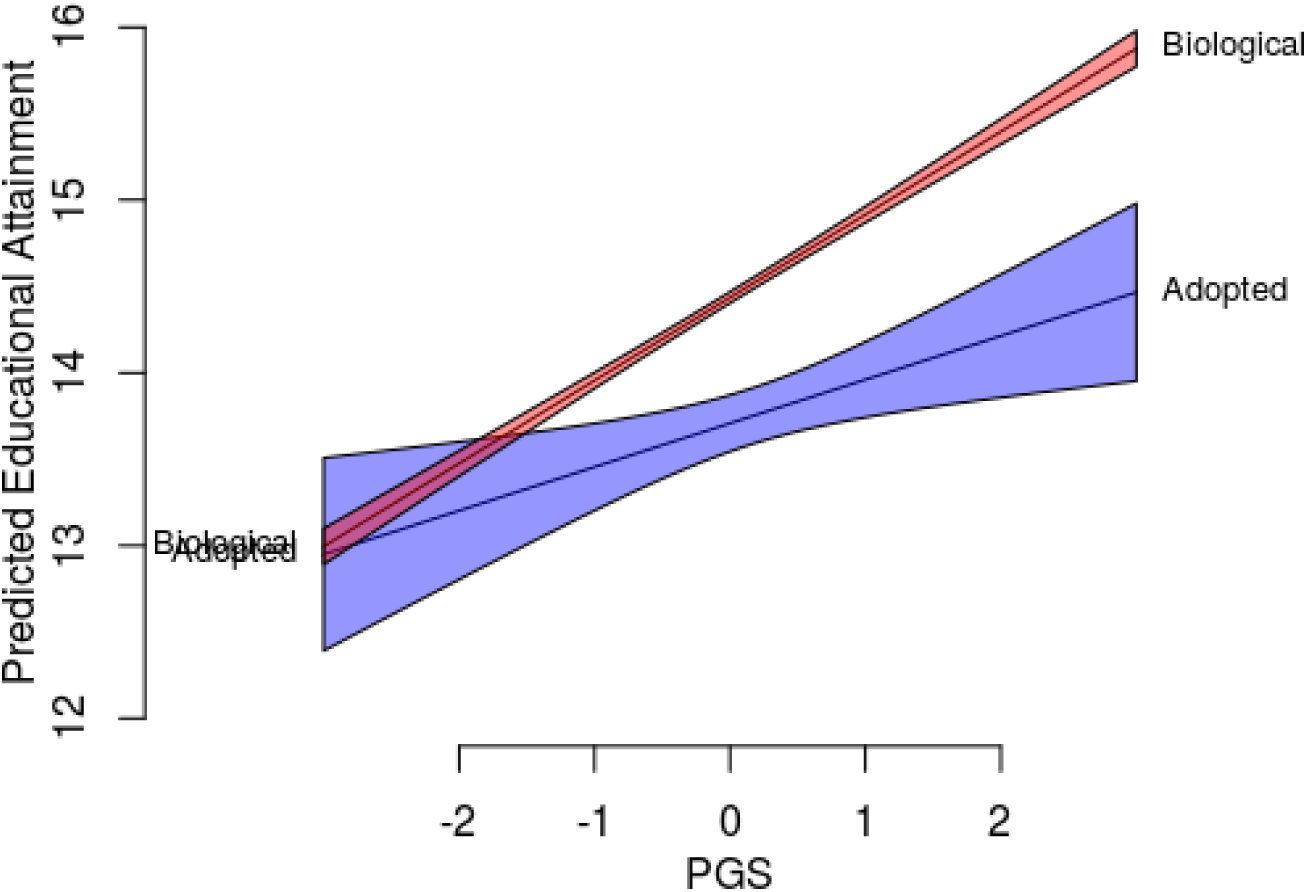
A comparison of predicted educational attainment as a function of parental PGS and child status (biological or adopted).

One issue with these comparisons between families who adopt and those who do not is our finding that parental PGS is higher among adoptive parents (Table 1), which suggests the possibility of other family-level differences between families with adopted versus biological children. As a sensitivity test, we also conducted analysis using only those families who have both adopted and biological children (44). Table 3 reports results for families with both adoptive and biological children. Our baseline results continue to support the idea that parental PGS predicts adoptee educational attainment (although, we note the result becomes insignificant in the saturated model). Note that even in these families with both biological and adopted children, we also find that biological children complete nearly three quarters of a year of schooling more on average compared with adopted children.

**Table 3.** Associations between child education and parental polygenic score in families with both adoptive and biological children.

|  | Baseline | Saturated |
| --- | --- | --- |
| Intercept | 13.708 | 14.140 |
| SE | 0.105 | 0.156 |
| p-value | <1E-5 | <1E-5 |
| PGS | 0.253 | 0.229 |
| SE | 0.113 | 0.134 |
| p-value | 2.57E-02 | 8.90E-02 |
| Bio Child | 0.733 | 0.700 |
| SE | 0.140 | 0.207 |
| p-value | <1E-5 | 7.49E-04 |
| PGS:Bio Child | 0.296 | 0.292 |
| SE | 0.154 | 0.151 |
| p-value | 5.41E-02 | 5.39E-02 |
| Male Child |  | -0.085 |
| SE |  | 0.209 |
| p-value |  | 6.84E-01 |
| Child Birthyear |  | -0.097 |
| SE |  | 0.023 |
| p-value |  | 2.34E-05 |
| Bio Child:Male Child |  | -0.541 |
| SE |  | 0.270 |
| p-value |  | 4.50E-02 |
| PGS:Male Child |  | -0.013 |
| SE |  | 0.173 |
| p-value |  | 9.42E-01 |
| Bio Child:Child Birthyear |  | 0.087 |
| SE |  | 0.035 |
| p-value |  | 1.36E-02 |
| PGS:Child Birthyear |  | 0.014 |
| SE |  | 0.018 |
| p-value |  | 4.27E-01 |
| N | 1491 | 1486 |

## 4. Discussion

A large and growing literature leverages the rapid and important advances in genotyping and prediction using genetics and the associated creation of polygenic scores to test novel hypotheses about the links between genetics and phenotypes of interest. A key component of this literature focuses on estimating statistical associations between individual genotype and phenotype with a parsimonious set of controls based on the reasoning that genetic variation precedes all developmental processes and is thus independent of environmental feedback.

A limitation of this approach is that, due to the nature of meiosis, a child’s genotype is highly correlated with the genotype of their biological parent. By extension, the child’s genotype may also be correlated with environmental inputs during childhood (to the extent that the parental genotype is correlated with such inputs). Children with a high PGS for education are likely to have grown up in households with highly educated parents and both influences—that of the child’s genetics and that of their environments—likely contribute to the children’s educational attainments. Children with high PGS for education will experience differences in their surrounding environments—neighborhoods (23,45), for example—compared to children with a lower PGS.

In an attempt to separate children’s genetic effects from parental effects, earlier studies (31,32) used the fact that half of each parent’s alleles are not transmitted to children. If GWAS findings related to educational attainment were due to entirely direct (i.e., within-person) genetic effects, then the non transmitted alleles should contain no predictive information about the biological offspring’s educational attainment, net of the directly transmitted alleles. Yet this was not the case; a sizeable portion of the “genetic effect” on children’s schooling was estimated to be due to indirect genetic effects. In contrast, parallel findings for other phenotypes suggested a much smaller role for indirect genetic effects (31).

Our work is motivated by a different study design meant to capture a broader measure of parental effects. Building on a tradition of earlier work in both behavior genetics (46) and economics (47–49) that used adopted children to separate child genotype and parental genotype/environment, we test whether the polygenic score of parents predicts the educational attainment of their adopted offspring. We utilize the fact that, given the absence of direct genetic inheritance, there is no mechanical (i.e. transmitted) correlation between the polygenic score of a parent and adopted child. We find that a one-SD increase in the PGS of a parent predicts roughly 0.25 years of schooling for an adopted offspring as compared to roughly 0.5 years of schooling for a biological offspring. Our findings are consistent with those that utilized the design based on non-transmitted alleles (31,32) in suggesting that the educational attainment polygenic score is capturing both direct and indirect genetic (i.e. parental) effects.

While our results are the first to use our proposed research design, we acknowledge several limitations. The WLS is representative of the high school graduating class of 1957 in Wisconsin and a selected sibling. While Wisconsin is broadly representative of the US during this period (36), we acknowledge that our results may not be generalizable to other populations or cohorts. This limitation would be larger to the extent that individuals in Wisconsin during this period had different adoption patterns than individuals not represented in this sample. This limitation is countered by the recognition that very few datasets contain the information we require for our analysis, including two generations of family members, information on adoptive status and completed schooling in the children’s generation, and genetic information. A second limitation is that we have incomplete information about the adoption process for the families in our data, including a lack of information about selectivity in the adoption process and at what age the adoptee joined the adoptive family. Results from Table 3, wherein we focus only on families that choose to adopt, are meant to ensure that our results are not explained by selection along these lines. To the extent that adopted children begin living with their adoptive parents at later ages in our sample (compared with at birth), this would likely lead our estimates to be relatively conservative estimates of the association between parental PGS on the adopted children’s outcomes.

Our findings suggest the standard approach to linking genotype-phenotype using polygenic scores may inadvertently include the influence of both indirect and direct genetic effects. An implication of our results, that parental genotype is associated with *adopted* children’s phenotypes, is that estimates of the associations between own-genotype on own-phenotypes are not strictly capturing direct genetic effects but instead are confounded by family background. The magnitude of this confounding—labeled “genetic nurture” in recent studies— may well vary by phenotype. Educational attainment is a phenotype with a relatively large amount of confounding by genetic nurture (31).

While other studies have produced evidence of the existence of confounding of polygenic scores, the data requirements are restrictive. For example, Kong et al. (31) required trio data from families in the Icelandic population to generate evidence of genetic nurture. Our results add a new test of this confounding by making use of adoptive children and parental PGS data. While each approach has limitations, we think that the consistent findings from both approaches are instructive. Our results thus add to the accumulating evidence that PGS effects should be interpreted with caution and are not pure “genetic” effects for educational attainment and likely other phenotypes. This suggests that since polygenic scores, as currently constructed, combine genetic and environmental influences, new research designs—perhaps including the use of within family studies at the GWAS stage (50) or attempts to estimate structural parameters associated with genetic nurture (26)—will need to be used to separate these estimated impacts.

## Acknowledgements

This research uses data from the Wisconsin Longitudinal Study (WLS) of the University of Wisconsin-Madison. Since 1991, the WLS has been supported principally by the National Institute on Aging (AG-9775, AG-21079, AG-033285, and AG-041868), with additional support from the Vilas Estate Trust, the National Science Foundation, the Spencer Foundation, and the Graduate School of the University of Wisconsin-Madison. Since 1992, data have been collected by the University of Wisconsin Survey Center. A public use file of data from the Wisconsin Longitudinal Study is available from the Wisconsin Longitudinal Study, University of Wisconsin-Madison, 1180 Observatory Drive, Madison, Wisconsin 53706 and at http://www.ssc.wisc.edu/wlsresearch/data/. The opinions expressed herein are those of the authors. This research was supported by core grants to the Center for Demography and Ecology at the University of Wisconsin-Madison (P2C HD047873) and to the Center for Demography of Health and Aging at the University of Wisconsin-Madison (P30 AG017266) and by award 96-17-04 from the Russell Sage Foundation and the Ford Foundation. The authors thank Ron Lee and participants at the Population Association of America 2018 conference for helpful suggestions.

## Compliance with Ethical Standards

### Informed consent

Informed consent was obtained from all individual participants included in the study.

### Ethical Approval

All procedures performed in studies involving human participants were in accordance with the ethical standards of the institutional and/or national research committee (Stanford IRB #39751 & University of Wisconsin Madison Minimal Risk IRB 2013-1477) and with the 1964 Helsinki declaration and its later amendments or comparable ethical standards

